# Charting the diversity of Uncultured Viruses of Archaea and Bacteria

**DOI:** 10.1101/480491

**Authors:** Fh Coutinho, F Rodríguez-Valera

**Affiliations:** Evolutionary Genomics Group, Departamento de Produccíon Vegetal y Microbiología, Universidad Miguel Hernández, Campus San Juan, San Juan, Alicante 03550, Spain

## Abstract

Viruses of Archaea and Bacteria are among the most abundant and diverse biological entities on Earth. Unraveling their biodiversity has been challenging due to methodological limitations. Recent advances in culture-independent techniques, such as metagenomics, shed light on viral dark matter, revealing thousands of new viral genomes at an unprecedented scale. However, these novel genomes have not been properly classified and the evolutionary associations between them were not resolved. Here, we performed phylogenomic analysis of nearly 200,000 viral genomic sequences to establish GL-UVAB: Genomic Lineages of Uncultured Viruses of Archaea and Bacteria. GL-UVAB yielded a 44-fold increase in the amount of classified genomes. The pan-genome content of the identified lineages revealed their infection strategies, potential to modulate host physiology and mechanisms to escape resistance systems. Furthermore, using GL-UVAB for annotating metagenomes from multiple ecosystems revealed elusive habitat distribution patterns of viral communities. These findings expand the understanding of the diversity, evolution and ecology of viruses of prokaryotes.

## Introduction

Grasping the biodiversity of viruses of Bacteria and Archaea has been a major challenge within the field of virology. Limitations for viral cultivation and purification associated with the absence of universal marker genes have been major drawbacks in the effort to chart and classify the biodiversity of these viruses[1,2]. The taxonomic classification system established for viruses of Bacteria and Archaea was originally based on morphological traits, but genetic studies demonstrated that the major taxa established through this approach are not monophyletic[3–5]. Thus, viral classification and taxonomy has come to rely heavily on comparative genomics. This shift has led the International Committee for the Taxonomy of Viruses (ICTV) to call for a scalable genome-based classification systems that can also be applied to uncultured viruses for which no phenotypic data is available[6].

Phylogenomic trees and genomic similarity networks incorporate full genomic data for comparison and clustering of viral genomes. Both phylogenomic and network based approaches have showed promising results for reconstructing phylogenies and classifying and identifying novel viral taxa[1,5,7]. These approaches circumvent the biases and limitations associated with morphological data or the use of phylogenetic markers, and are easily scalable to thousands of genomes[5,8]. Network methods rely on the identification of orthologous groups shared among genomes, which can be problematic for viruses due to the ratio in which their genes evolve. Additionally, the evolutionary associations among genome clusters identified by network approaches are not explicitly resolved by these methods[5,9]. Meanwhile, phylogenomic approaches provide trees in which the associations among genomes are easily interpretable under an evolutionary perspective. For these reasons, phylogenomic methods have been the standard approach for reconstructing phylogenies of prokaryotic viruses[1,4,7,8,10–14]. Previous studies have leveraged this method to investigate the genetic diversity of cultured viruses, but overlooked the uncultured diversity.

Thousands of novel viral genomes were recently discovered through culture-independent approaches, such as shotgun metagenomics, fosmid libraries, single-virus sequencing and prophage mining[4,11,15–18]. These new datasets unraveled an extensive biodiversity that had been overlooked by culture-based approaches. These genomes have the potential to fill many of the gaps in our understanding of the diversity of viruses of prokaryotes. Yet, achieving this goal requires that these genomes are properly organized in a robust evolutionary framework. Here, we applied a phylogenomic approach to chart the diversity of uncultured viruses of Bacteria and Archaea aiming to gain insights on their genetic diversity, evolution and ecology.

## Results

### Phylogenomic reconstruction

A database of genomic sequences of prokaryotic viruses was compiled with isolated dsDNA viruses from RefSeq and uncultured viruses that were discovered across multiple ecosystems using approaches that bypassed culturing. This database amounted to 195,698 viral genomic sequences along with associated information of computational host predictions and ecosystem source (Table S1). Following pre-filtering and redundancy removal steps, a subset of 16,484 genomic sequences was selected for phylogenomic reconstruction (Methods and Figure S1). An all-versus-all comparison of the protein sequences encoded in this dataset was performed and used to calculate Dice distances between genomes. Essentially, the Dice distances between a pair of genomes decreases the more proteins that are shared between them and the higher their degree of identity. Finally, the obtained matrix of Dice distances was used to construct a phylogenomic tree through neighbor-joining (Figure 1). In addition, a benchmarking dataset containing a subset of 2,069 dsDNA prokaryotic viral genomes from NCBI RefSeq was analyzed in parallel for comparison and validation of results. The robustness of the tree topology was evaluated through a sub-sampling approach. Nodes displayed high confidence values (average 50% recovery), and 90% of all nodes were recovered at least once among the re-sampled trees. These figures were obtained when reducing the data used to calculate distances to approximately 64% of the amount used to establish the original tree, demonstrating that tree topology is robust even in the presence of incomplete or fragmented genomes, which might be the case for some of the uncultured viral genomes used.

**Figure 1.**
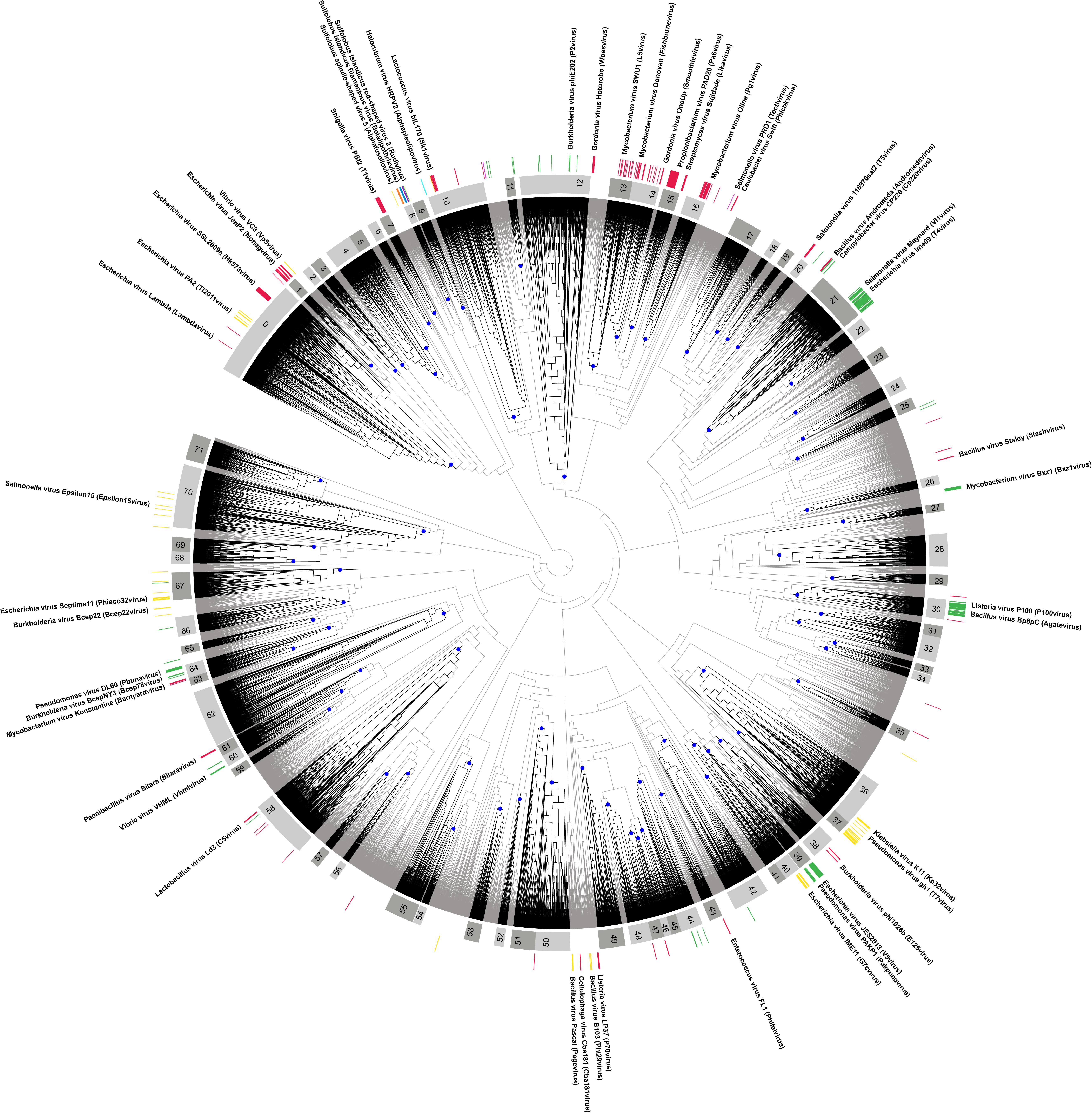
Phylogenomic reconstruction of 16,484 viral genomic sequences reveals major lineages of uncultured prokaryotic viruses. The tree was built through Neighbor-Joining based on Dice distances calculated between viral genomic sequences from both NCBI RefSeq and those reconstructed from metagenomes, fosmid libraries, single virus genomes and prophages integrated into prokaryote genomes. Tree was midpoint rooted. Branch lengths were omitted to better display tree topology. Each of the 72 Level-1 GL-UVAB lineages are highlighted by black colored branches and with their defining nodes in indicated by blue dots. Numeric identifiers for the lineages are displayed in the innermost ring within gray strips. The outermost ring depicts the ICTV family level classification assignments of RefSeq viral genomes that were included in the tree. Color codes for ICTV families are as follows: Myoviridae (Green), Siphoviridae (Red), Podoviridae (Yellow), Lipothrixviridae (Blue), Fuselloviridae (orange), Sphaerolipoviridae (Cyan), Rudiviridae (Purple) and Tectiviridae (Pink). Fore reference, representatives of isolated viral genomes are shown with their ICTV genus level classification shown in parentheses.

### Clustering uncultured prokaryotic viruses into closely related lineages

A three-step approach was applied to categorize diversity into hierarchical levels of increasing genomic relatedness: Level-1 (average Dice distances between genomes equal or below 0.925, and number of representatives equal or above 50), Level-2 (average Dice distances between genomes equal or below 0.825, and number of representatives equal or above 10) and Level-3 (average Dice distances between genomes equal or below 0.5, and number of representatives equal or above 3). These cutoffs were selected to establish lineages at a degree of genomic similarity within the range observed for families, sub-families and genera established by the ICTV (Table S4). At the first level, 10,357 genomic sequences were assigned to 72 lineages (Figure 1). Of these, 68 of the Level-1 lineages have at least one complete viral genome assigned to them. In addition, 50 of these Level-1 lineages had at least one reference viral genome assigned to them. At the second level, 10,356 sequences were assigned to 314 lineages, while at the third level, 6,971 sequences were assigned to 942 lineages. This three-level classification system was used to establish the GL-UVAB. Apart from 6 cases, all of the Level-1 lineages are composed of genomes assigned to a single taxonomic family as defined by the ICTV. Out of the 942 Level-3 lineages, 104 of them included genomes classified at the level of genus by the ICTV. Of these, 74 lineages are consistent regarding the genus, meaning that all classified genomes are assigned to the same genera. Meanwhile, only 6 genera were split among more than one Level-3 lineage. Thus, the identified lineages of uncultured viruses are in agreement with the ICTV established taxonomy.

Genome sequences that were not included in the phylogenomic reconstruction were assigned to the lineage of their closest relatives as determined by the average amino acid identity and percentage of shared genes. Following this lineage expansion step, a total of 100,907 sequences were classified to at least one level (Table S1), which represents a 44-fold improvement in the proportion of classified sequences compared to the amount of RefSeq prokaryotic viral genomes classified by the NCBI taxonomy database. The dataset that contributed most (∼60%) of the classified sequences was from a cross-ecosystem global analysis of metagenomes[16], followed by metagenomes from the Tara and Malaspina expeditions [17] (∼15%), global marine viromes[15] (∼10%) and prophages identified in bacterial genomes[18] (∼10%) (Figure S3).

### Targeted hosts and ecosystem sources of GL-UVAB lineages

GL-UVAB lineages differed regarding host prevalence (Figure 2A). Out of the 72 Level-1 lineages, 34 are predicted to infect a single host phylum, most often *Proteobacteria*, *Firmicutes* or *Actinobacteria*, while 35 lineages are predicted to infect two or more phyla. Level-3 lineages display the highest levels of host consistency. Among Level-3 lineages with at least one annotated host, 99% of them are predicted to infect a single phylum and 76% are predicted to infect a single genus. Lineages also differed regarding the ecosystem sources from where their members were obtained (Figure 2B). Nearly all lineages contained members obtained from multiple ecosystems but aquatic and human-associated samples were consistently the main sources of genomes, highlighting diversity within these habitats and the potential for discoveries in other under-explored ecosystems. Trends of host and ecosystem prevalence observed for the expanded lineages (Figure S4) were consistent with those obtained from the analysis of the original dataset, corroborating the validity of these patterns. Applying the same lineage identification criteria to the benchmarking dataset of RefSeq viral genomes identified 12 Level-1 lineages, 49 Level-2 lineages and 162 Level-3 lineages (Figure S2). Among these Level-3 lineages, 145 (91%) are composed of genomes that infect within the same host order. This observation further corroborates the validity of the associations between GL-UVAB lineages and targeted hosts.

**Figure 2.**
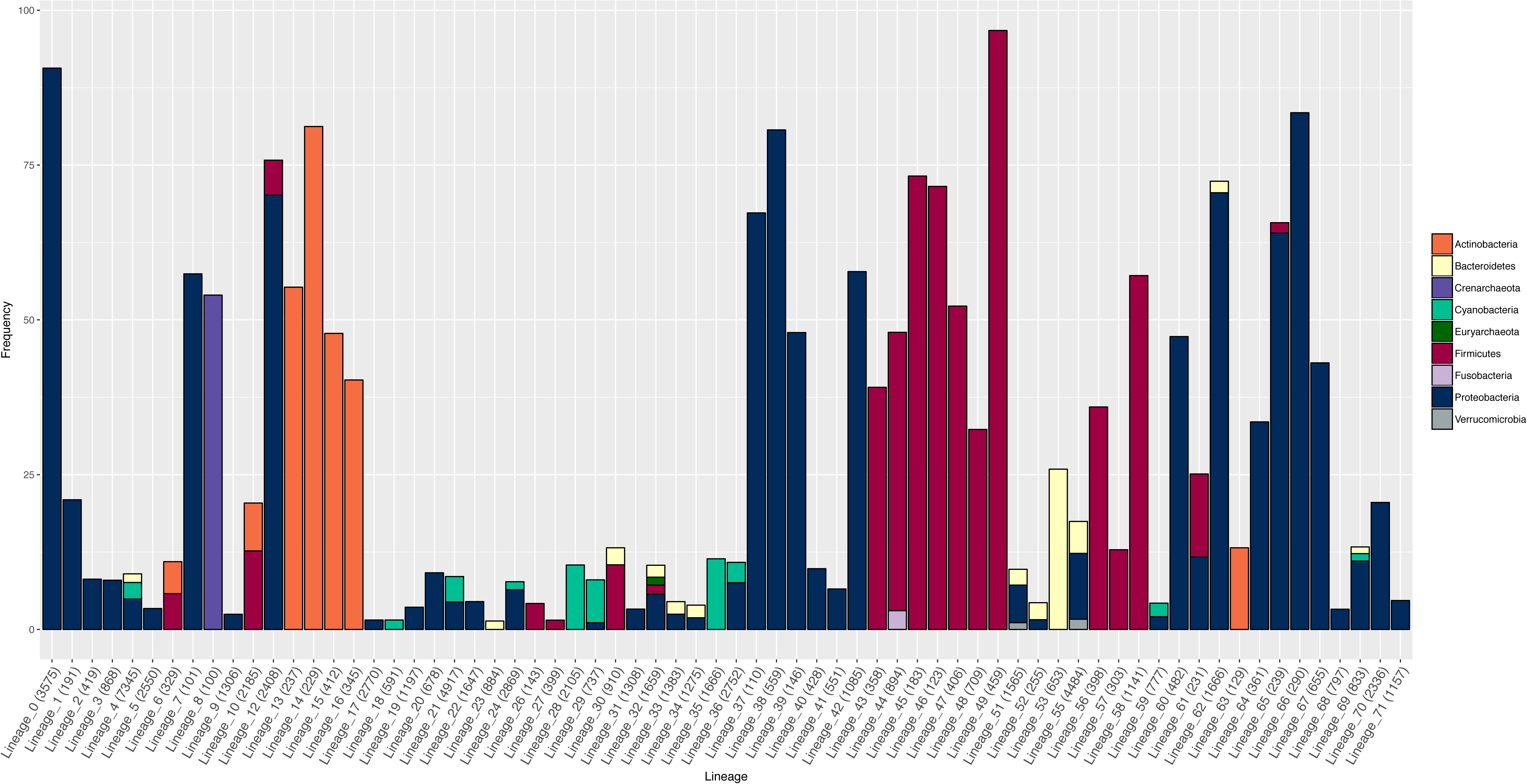
Prevalence of targeted host and ecosystem sources among Level-1 GL-UVAB lineages assigned through phylogenomic reconstruction. A) Frequency of infected host phyla across each of the 72 identified lineages. B) Frequency of ecosystem sources from which viral sequences were obtained across each of the 72 identified lineages. For clarity, only hosts and ecosystems with prevalence equal or above 1% are shown. Numbers in parentheses indicate the total number of genomes assigned to each lineage.

### GL-UVAB lineages differ in habitat distribution and pan-genome content

The observed differences in host preference and ecosystem source among lineages led us to investigate the applicability of GL-UVAB as a reference database for deriving abundance profiles from metagenomes. We analyzed the abundances of 72 GL-UVAB Level-1 lineages across metagenomes from marine, freshwater, soil and human gut samples (Figure 3). Lineages 28, 21, and 36 were the most abundant in marine samples, in agreement with the high prevalence of *Cyanobacteria* and *Proteobacteria* as hosts of these lineages (Figure 2A). Meanwhile, the lineages 64 (which mostly infects *Proteobacteria* of classes Gamma and Delta), 23 (*Proteobacteria* and *Bacteroidetes*), and 21 where the dominant groups in freshwater habitats. In temperate soil samples, the most abundant lineages were 14 (*Actinobacteria*), 38 (*Beta- and Gamma-Proteobacteria*) and 62 (*Gamma*p*roteobacteria*). Finally, human gut samples were dominated by lineages 64, 30 (*Firmicutes and Bacteroidetes*) and 10 (*Firmicutes* and *Actinobacteria*).

**Figure 3.**
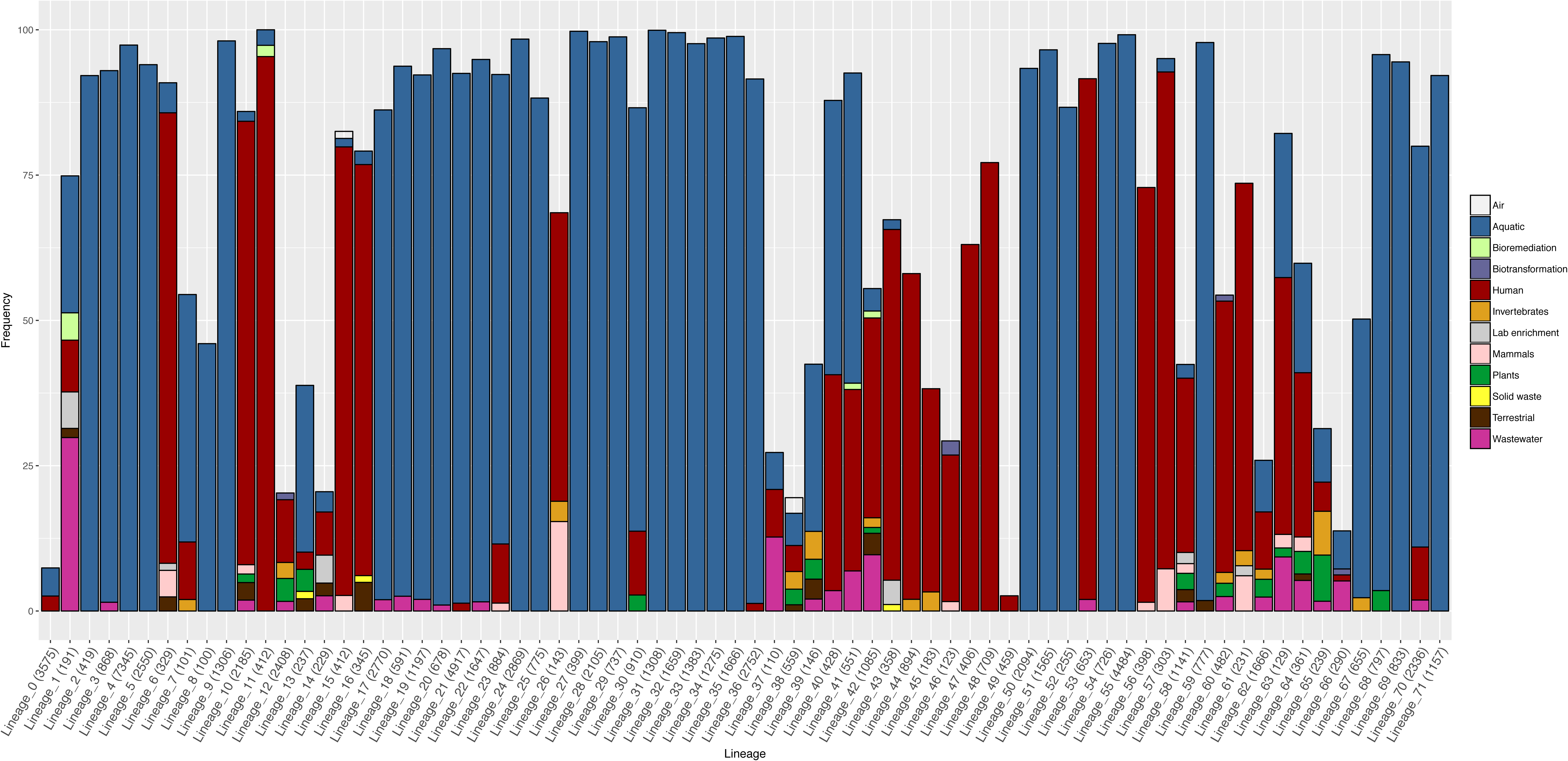
Abundance patterns of GL-UVAB Level-1 lineages across habitats. Bar plots display the average and standard errors of the relative abundances of GL-UVAB Level-1 lineages across metagenomes and metaviromes from marine, freshwater, human gut and soil ecosystems.

We inspected the pan-genome of the identified lineages by clustering their protein encoding genes into orthologous groups (OGs). A total of 79,281 OGs containing at least three proteins were identified. These OGs displayed a sparse distribution, i.e., were only detected in a small fraction of genomes within lineages (Table S3). The most conserved OGs encoded functions associated with nucleic acid metabolism and viral particle assembly. Few OGs encoded putative auxiliary metabolic genes (AMGs), and those where never shared by all the members of a lineage. A total of 1,755 promiscuous OGs, present in the pan-genome of three or more Level-1 lineages were identified.

## Discussion

Only a small fraction of prokaryotic viruses can be cultivated through currently available laboratory techniques. This limitation has left many gaps in our understanding of their biodiversity. The results presented here help to bridge these gaps by leveraging on a large dataset of viral genomic sequences obtained without cultivation from multiple ecosystems. Categorizing this new diversity into a robust evolutionary framework brings us closer to properly grasping the extension of viral biodiversity.

The average dice distance cutoffs used for defining lineages were chosen to classify as many genomes as possible while maintaining cohesiveness within lineages regarding similarity between genomes, targeted hosts and taxonomic classification as defined by the ICTV. These goals were achieved, as the GL-UVAB lineages are formed by groups of closely related genomes (Table S4), which was reflected in their targeted hosts (Figure 2A), pan-genome content (Table S2) and taxonomic classification (Table S5). Using different cutoffs for lineage identification would have resulted in the distinct lineages and consequently different patterns of host and source prevalence, pan-genome content, and habitat distribution. Nevertheless, GL-UVAB was conceived to be an evolving system. We encourage researchers to adapt the GL-UVAB approach to suit the needs of the specific questions under investigation. For example, performing species-level clustering genomes would require average Dice distance cutoffs even lower than those used to delineate Level-3 lineages.

The re-sampling approach showed that the tree topology was robust even in the presence of fragmented genomes. Still, incomplete genomes are more likely to be misclassified by our pipeline. Yet, in the absence of complete genomes, these fragments are the best available option for obtaining at least a provisional classification of uncultured viruses. As the quality and amount of uncultured viral genomes discovered increases, new data can be used to update and improve the GL-UVAB.

The consistency of the targeted hosts among lineages identified with our phylogenomic approach suggests that the assignment to GL-UVAB lineages provides a rough estimate of the hosts of uncultured viruses. This is of fundamental importance, considering the growing diversity of viral genomes discovered from metagenomic datasets for which no host information is initially available[19,20]. Host prevalence analysis indicated that approximately half of the Level-1 lineages are capable of infecting more than a single host phylum (Figure 2A). The ability to interact with the molecular machinery of the host is a major driver of the evolution of prokaryotic viruses. Thus, closely related genomes (that belong to the same lineages) likely have undergone similar evolutionary pressures that ensure host infectivity, leading to the observed pattern of higher host consistency among the lowest level of hierarchical classification, i.e., Level-3 lineages. Meanwhile, the ability of some lineages to infect across multiple host phyla is likely an indication of the high level of genomic plasticity of viruses that allows them to evolve to infect new organisms that are not closely related to their original hosts.

Metagenomic studies of viral ecology strive to elucidate the habitat distribution patterns of taxa across ecosystems. The abundance patterns observed for the GL-UVAB lineages (Figure 3) are a reflection of their distinctive trends of host prevalence (Figure 2A). As expected, the GL-UVAB lineages that dominated at each ecosystem often targeted taxa that are the most abundant at these habitats[21,22], e.g., lineages that target *Proteobacteria* and *Cyanobacteria* at aquatic samples and lineages that target *Bacteroidetes* and *Firmicutes* in the human gut. Although this observation might seem obvious, it does not emerge when using cultured viral genomes for the taxonomic annotation of metagenomes. Instead, the same taxa are often observed with similar abundance patterns regardless of the ecosystem sampled. This occurs because established taxa have no discernible host or ecosystem preferences and because much of viral diversity is not encompassed by viral taxonomy[14,23,24]. Thus, the cohesiveness of GL-UVAB lineages regarding phylogeny, host preference and ecology allows for meaningful habitat-taxa associations to be observed.

A detailed investigation of the pan-genome content of the Level-1 lineage 21, revealed some of the strategies applied by these viruses during infection. This lineage was among the dominant group in both freshwater and marine samples and infects *Cyanobacteria* and *Proteobacteria.* The pan-genome of lineage 21 includes OGs encoding high-light inducible proteins, photosystem II D1 proteins and a transaldolase. These proteins are involved in photosynthesis and carbon fixation pathways[25]. Therefore, the success of this group across aquatic ecosystems might be linked to their capacity to use such proteins as AMGs to modulate the metabolism of their Cyanobacterial hosts during infection, redirecting it to the synthesis of building blocks to be used for the assembly of novel viral particles[25].

The promiscuous distribution observed for multiple OGs could be the result of the positive selection of these genes following events of horizontal gene transfer. Indeed, promiscuous OGs often encoded proteins that might confer advantages during infection. Five of them encode thymidylate synthase, a protein involved in nucleotide synthesis. Meanwhile, three promiscuous OGs encode the *PhoH* protein, which mediates phosphorus acquisition in nutrient deprived conditions. These findings suggest a selective pressure favoring the acquisition of genes that allow viruses to modulate host metabolism towards the production of nucleic acids to be used for the synthesis of progeny DNA[25]. Nevertheless, the sparse distribution of these and other auxiliary metabolic genes across lineages suggest that such genes have been recently acquired by the members of each lineage through horizontal gene transfer (HGT), rather than being present in the common ancestor of the group. Multiple methylases were identified among promiscuous OGs. Viruses use these proteins to protect their DNA from host restriction modification systems[26]. Prokaryotes can acquire restriction modification systems through HGT[27], and our data suggest that viruses also benefit from HGT by acquiring novel methylases that allow them to escape these systems. Finally, lysins (e.g., peptidases and amidases) were a common function among promiscuous OGs. This finding is surprising because lysins are believed to be fine-tuned for the specific structure of host cell wall[28,29]. Acquisition of novel lysins might help viruses to expand their host spectra or as a mechanism to ensure infectivity following the emergence of resistance mutations that lead to alterations in the structure of the host cell wall.

In conclusion, by analyzing thousands of uncultured viral genomes we were able to categorize the diversity of these biological entities. This was achieved by identifying lineages of uncultured viruses through a robust and scalable phylogenomic approach. Analyzing host and source prevalence, pan-genome content and abundance in metagenomes painted a more accurate picture of viral biodiversity across ecosystems. As these uncultured viruses are isolated and their morphology and host spectra are elucidated, they will be properly integrated into the ICTV classification system. Here we provided an initial framework for their taxonomic classification as well as insights regarding their ecology and genetic diversity. Future studies will continue to shed light on the viral dark matter across our planet’s many ecosystems. Our work provides the initial steps for a genome-based classification of these yet undiscovered evolutionary lineages, providing a solid framework to investigate the biology of prokaryotic viruses.

## Methods

### Viral genome database

The NCBI RefSeq dataset was used as a starting set of reference viral genome sequences. Host information for these sequences was retrieved from GenBank files, and their taxonomic classification was obtained both from the NCBI Taxonomy database and from the ICTV [30]. Additionally, genomic sequences were compiled from studies that used high throughput approaches to obtain viral genomes through culture-independent analysis, along with their associated host and ecosystem source information whenever available. These genomes were obtained from: environmental metagenomes and metaviromes[3,12,15–17,31,32], fosmid libraries of Mediterranean viruses[4,11], single virus genomes[33] and prophages integrated into prokaryotic genomes[18]. This dataset (henceforth referred to as Vir_DB_Nuc) contained a total of 195,698 viral genome sequences (Table S1 and Supplementary file 1). Protein encoding genes (PEGs) were predicted from Vir_DB_Nuc using the metagenomic mode of Prodigal[34], which identified 4,332,223 protein sequences (henceforth referred to as Vir_DB_Prot, Supplementary file 2). The Vir_DB_Prot dataset was queried against the NCBI-nr protein database using Diamond[35] for taxonomic and functional annotation.

### Sequence pre-filtering

Identifying viral sequences within metagenomic and metaviromic datasets can be problematic. Because each study used different strategies to achieve that goal we pre-filtered genome sequences from Vir_DB_Nuc to ensure that only *bona fide* prokaryotic viral genomes were included in downstream analyses. First, the Vir_DB_Prot dataset was queried against the prokaryotic virus orthologous groups (pVOGs)[36] protein database using diamond[35] (more sensitive mode, BLOSUM45 matrix, identity ≥ 30%, bitscore ≥ 50, alignment length ≥ 30 amino acids and e-value ≤ 0.01). For each sequence, the percentage of proteins mapped to the pVOGs database and the added viral quotient (AVQ) were calculated (as the sum of the individual viral quotient of the best hit of each protein). Hits to pVOGs that presented homology with proteins from Eukaryotic viruses were not considered. Sequences with 20% or more of the proteins mapped to the pVOGs database and with an AVQ equal to or greater than 5 were classified as *bona fide* prokaryotic viral genomes. This initial round of recruitment yielded 26,610 genomic sequences (Vir_DB_Nuc_R1). Next, proteins from the Vir_DB_Nuc_R1 dataset were used as bait for a second recruitment round. The remaining protein sequences (which were not recruited in the first round) were queried against Vir_DB_Nuc_R1 through diamond as described above. Genomic sequences from which at least 20% of the derived proteins mapped to a single genome from Vir_DB_Nuc_R1, yielding a minimum of three hits, were recruited to Vir_DB_Nuc_R2 (78,295 genomic sequences). Finally, a last step of manual curation was performed, which recruited mostly long contigs with high AVQ that did not match the percentage criteria of the automatic recruiting steps due to their high number of encoded proteins. This step recruited a total of 6,421 genomic sequences (Vir_DB_Nuc_R3).

We benchmarked the accuracy of the automatic recruiting steps with two datasets. First, a subset of Vir_DB_Nuc comprised of all the reference viral genomes was run through the recruitment pipeline using the same criteria described above. None of the 7,036 eukaryotic viruses were recruited by the pipeline (i.e., 100% precision) and 2,136 out of 2,297 prokaryotic viruses were correctly recruited (i.e., 92.99% recall). We also benchmarked the filtering pipeline with a dataset of 897 Gbp of genome sequence data derived from the NCBI RefSeq prokaryote genomes spanning 880 genera from 35 phyla. Sequences were split into fragments of 5, 10, 15, 20, 25, 50 and 100 Kbp to mimic metagenomic scaffolds. Using the filtering criteria described above and the subsequent length filtering for sequences longer than 30 Kbp would recruit only 109 sequences (0.36%), all of which displayed homology to the prophage sequences described by Roux et al. [18].

### Phylogenomic reconstruction

Phylogenomic reconstruction was performed using a subset of genomes from Vir_DB_Nuc that included all dsDNA RefSeq viral genomes annotated as complete for which the host Domain was either Bacteria or Archaea and the uncultured *bona fide* prokaryotic viruses from Vir_DB_Nuc_R3 with a length equal or greater than 30 Kbp or annotated as complete and with a length equal or greater than 10 Kbp. These criteria were established to ensure any incomplete genomes fragments encompassed a significant fraction of the complete genome (considering the 47 Kbp median size of complete genomes from RefSeq viruses of Bacteria and Archaea). Genome sequences were clustered with CD-HIT[37] using a cut-off of 95% nucleotide identity and minimum 50% coverage of the shorter sequence to remove redundant sequences. The non-redundant dataset contained 16,484 viral genomic sequences that were used for phylogenomic reconstruction (Vir_DB_Phy). Distances between genomes were calculated based on a modified version of the Dice method[4]. First, an all-versus-all comparison of the PEGs derived from the Vir_DB_Phy dataset was performed through diamond[35] (more sensitive mode, identity ≥ 30%, bitscore ≥ 30, alignment length ≥ 30 amino acids and e-value ≤ 0.01). Next, distances between genome sequences were calculated as follows: D_AB_ = 1 - (2^*^(AB) / (AA+BB)), where AB is the bitscore sum of all the valid hits of genome A against genome B, while AA and BB are the bitscore sum of all the valid hits of genome A against itself and of all the valid hits of genome B against itself, respectively. The more homologous proteins are shared between genomes A and B, and the higher the degree of identity between these homologous proteins, the closer to zero the value of *D*_*AB*_ will be. Non homologous proteins should produce no hits when comparing genomes A against B, but will hit with themselves when comparing A against A and B against B. Therefore, when estimating *D*_*AB*_, non homologous proteins are penalized, increasing the value of *D*_*AB*_. The obtained Dice distances matrix was used as input to build phylogenetic tree through Neighbor-Joining algorithm[38] implemented in the Phangorn package of R. The obtained tree was midpoint rooted.

### Tree topology validation by re-sampling

A re-sampling approach was applied to test the consistency of the tree topology. First, 20% of the proteins encoded in the genomes used to build the tree were randomly selected. Then, distances between genomes were re-calculated after excluding any hits from the all-versus-all search in which either the query or subject sequences were selected for exclusion, which removes approximately 36% of all of the original hits. Finally, the obtained distance matrix was used to construct a new tree. This process was repeated over one hundred iterations. Next, we measured the frequency in which the nodes from the original tree were present in the re-sampled trees.

### Lineage identification

Lineage identification was performed by parsing the phylogenetic tree to identify monophyletic clades that matched the established criteria for maximum average Dice distances between genomes, and for a minimum number of representatives. Lineages were identified in three steps, aimed at capturing diversity into levels of increasing genomic relatedness: Level-1 (average Dice distances between genomes equal or below 0.925, and number of representatives equal or above 50), Level-2 (average Dice distances between genomes equal or below 0.825, and number of representatives equal or above 10) and Level-3 (average Dice distances between genomes equal or below 0.5, and number of representatives equal or above 3). These cutoffs were selected based on the degree of genomic similarity observed for taxa of prokaryotic viruses described by the ICTV and NCBI Taxonomy databases (Table S4), with the goal of classifying as many sequences as possible while maximizing consistency within clades regarding pan-genome content and agreement with the currently accepted viral taxonomy. To trace the pan-genomes of the identified lineages the proteins derived from 16,484 genomes used for phylogenomic reconstruction were clustered into orthologous groups using the orthoMCL algorithm[39] implemented in the Get_Homologues pipeline[40]. The MCL inflation factor was set to 1 and all other parameters were set to default.

### Lineage expansion by closest relative identification

Sequences that did not pass the initial length and redundancy filters to be included in the phylogenomic tree were assigned to the lineages of their closest relatives. Closest relatives were defined as the genome with highest percentage of matched protein encoding genes (PEGs) as detected by Diamond searches. Potential ties were resolved by choosing the closest relative with the highest average amino acid identity (AAI) value. A minimum AAI of 50% and the percentage of matched PEGs of 50% was required for closest relative assignments.

### Lineage benchmarking and validation with RefSeq prokaryotic viruses

We sought to validate our strategy for lineage identification. To that end a dataset of 2,069 genome sequences of dsDNA viruses of Archaea and Bacteria from the NCBI RefSeq database was used for benchmarking. The steps for distance calculation, tree construction and lineage identification were performed exactly as described for the full dataset. Next, we calculated statistics of distances between taxa established by ICTV and NCBI (Table S4).

### Lineage abundance in metaviromes and metagenomes

The abundances of Vir_DB_Nuc sequences were estimated in viral metagenomes (viromes) from the following ecosystems: marine epipelagic samples[41], healthy human gut[42], freshwater lakes[43], and because no large scale viromes of mesophilic soils were available we used cellular metagenomes from this ecosystem[44,45]. Sequencing reads from these metagenomes and metaviromes were retrieved from the European Nucleotide Archive or NCBI Short Read Archive. Subsets of twenty million R1 reads from each sample were mapped to Vir_DB_Nuc using Bowtie2[46] using the sensitive-local alignment mode. Lineage abundances across samples were calculated by summing the relative abundances of individual genomes according to their assigned lineages.

## Supporting information

## Acknowledgements

FHC and FRV were supported by the grants “VIREVO” CGL2016-76273-P [AEI/FEDER, EU], (co-funded with FEDER funds); Acciones de dinamización “REDES DE EXCELENCIA” CONSOLIDER-CGL2015-71523-REDC from the Spanish Ministério de Economía, Industria y Competitividad and PROMETEO II/2014/012 “AQUAMET” from Generalitat Valenciana. FHC was supported by APOSTD/2018/186 Post-doctoral fellowship from Generalitat Valenciana.

## Author contributions

FHC conceived, designed and performed experiments and analyzed the data. FHC and FRV wrote the manuscript.

## Supplementary Figure legends

**Figure S1.**
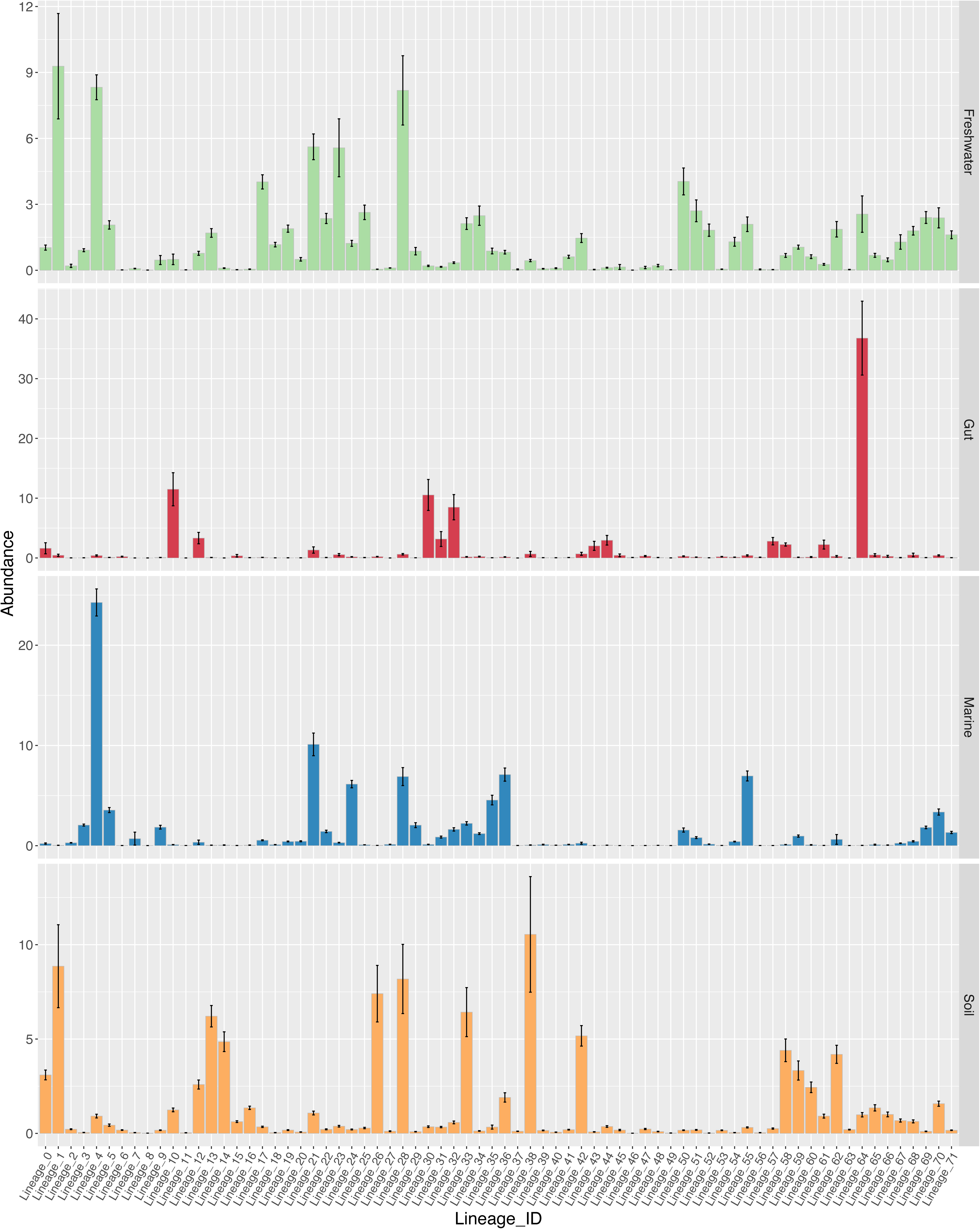
Flowchart summarizing the methodology used to establish GL-UVAB. The initial dataset of genomic sequences consisted of the NCBI RefSeq and viral genomic sequences obtained through culturing independent approaches adding up to 195,698 genomic sequences from which 4,332,223 protein encoding genes were identified. After the initial filtering, 16,484 sequences were selected for phylogenomic reconstruction. Dice distances were calculated between this set and the resulting distance matrix was used for phylogenomic reconstruction through Neighbour-Joining. The obtained tree was used to identify lineages at three levels, based on average Dice distances within nodes: Level-1 (average Dice distances between genomes equal or below 0.925, and number of representatives equal or above 50), Level-2 (average Dice distances between genomes equal or below 0.825, and number of representatives equal or above 10) and Level-3 (average Dice distances between genomes equal or below 0.5, and number of representatives equal or above 3). Lineage abundances were estimated in metagenomic datasets by read mapping. Lineage pan-genomes were determined by identifying clusters of orthologous genes. Finally, sequences that were not included in the original tree were assigned to the lineages by closest relative identification (CRI). Closest relatives were determined based on percentage of matched genes and average amino acid identity and setting a minimum value of 50% for both these variables.

**Figure S2.** Phylogenomic reconstruction of 16,484 viral genomic sequences. The tree was built through Neighbor-Joining based on Dice distances calculated between viral genomic sequences from both NCBI RefSeq and those reconstructed from metagenomes, fosmid libraries and prophages integrated into prokaryote genomes. The tree was midpoint rooted. Nodes were collapsed if all the leaves belonged to the same Level-1 lineage to better display higher-order associations between lineages.

**Figure S3.** Phylogenomic reconstruction of 2,069 genomes of dsDNA viruses of Archaea and Bacteria from RefSeq. The tree was built through Neighbor-Joining based on Dice distances calculated between complete prokaryotic viral genomes from NCBI RefSeq. The tree was midpoint rooted. Branches are colored according to their family level taxonomic classification, the inner ring displays classification of genomes into subfamilies and the outer ring displays classifications into lineages identified for the benchmarking dataset.

**Figure S4.** Dataset sources of GL-UVAB genomes. Bar plots depict the abundance of sequences from each original publication that made up the full dataset (Total), those that were recruited during the pre-filtering step (Recruited) and those that were classified into GL-UVAB lineages (Classified).

**Figure S5.** Prevalence of targeted host and ecosystem sources among Level-1 GL-UVAB lineages assigned through phylogenomic reconstruction and lineage expansion by closest relative identification. A) Frequency of infected host phyla across each of the 72 identified lineages. B) Frequency of ecosystem sources from which viral sequences were obtained across each of the 72 identified lineages. For clarity, only hosts and ecosystems with prevalence within a lineage equal or above 1% are shown. Numbers in parentheses indicate the total number of genomes assigned to each lineage.

## Supplementary files

**File S1.** Multifasta file containing the 195,698 viral genome sequences from Vir_DB_Nuc analyzed in this study.

**File S2.** Multifasta file containing the 4,332,223 protein encoding gene sequences predicted from Vir_DB_Nuc.

**File S3.** Newick format phylogenomic tree used to define GL-UVAB lineages. The tree was constructed with the Neighbor-Joining algorithm based on Dice distances calculated between 16,484 genomic sequences.

**Table S1.** Table containing detailed information of all the genomic sequences from Vir_DB_Nuc analyzed in this study, including sequence identifier, NCBI access number, original dataset, sequence length, number of identified PEGs, taxonomic classification (NCBI and ICTV), ecosystem source and host taxonomic classification and affiliation to the identified lineages at three hierarchical levels.

**Table S2.** Table describing the prevalence of OGs across GL-UVAB Level-1 lineages with associated taxonomic and functional annotation. Only OGs detected in at least 3 members of a lineage are shown.

**Table S3**. Statistics of the Dice distances observed between genomes of the taxa established by NCBI/ICTV and the lineages established by GL-UVAB.

**Table S4.** Prevalence of targeted hosts, ecosystem and dataset sources, taxonomic classification and completeness of genomes among the three levels of GL-UVAB lineages.

