## Supplementary material for "Charting the diversity of Uncultured Viruses of Archaea and Bacteria"

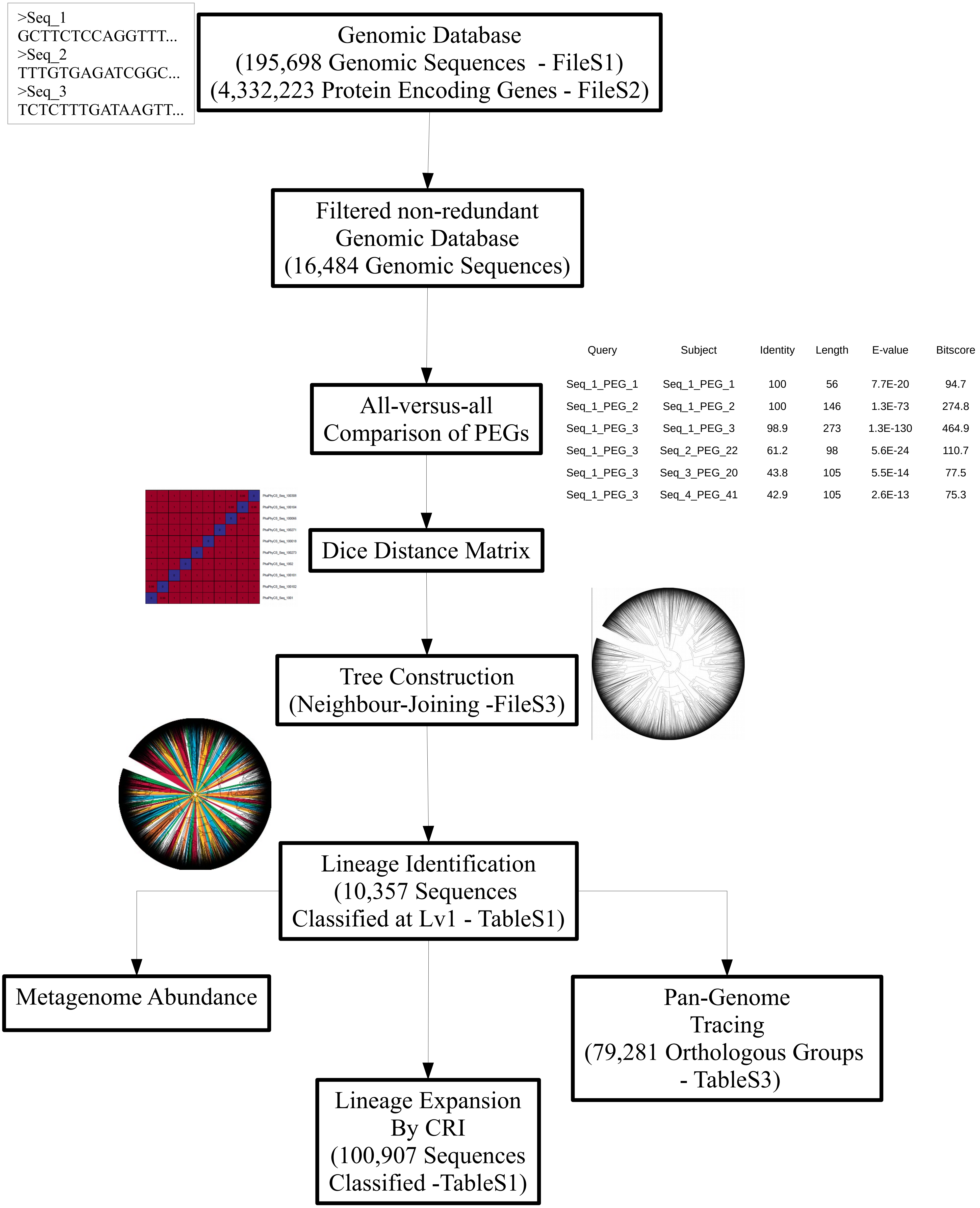


| Query | Subject | Identity | Length | E-value | Bitscore |
|---|---|---|---|---|---|
| Seq_1_PEG_1 | Seq_1_PEG_1 | 100 | 56 | 7.7E-20 | 94.7 |
| Seq_1_PEG_2 | Seq_1_PEG_2 | 100 | 146 | 1.3E-73 | 274.8 |
| Seq_1_PEG_3 | Seq_1_PEG_3 | 98.9 | 273 | 1.3E-130 | 464.9 |
| Seq_1_PEG_3 | Seq_2_PEG_22 | 61.2 | 98 | 5.6E-24 | 110.7 |
| Seq_1_PEG_3 | Seq_3_PEG_20 | 43.8 | 105 | 5.5E-14 | 77.5 |
| Seq_1_PEG_3 | Seq_4_PEG_41 | 42.9 | 105 | 2.6E-13 | 75.3 |
