## Supplementary figures and images for "Charting the diversity of Uncultured Viruses of Archaea and Bacteria"

### Supplementary file 8

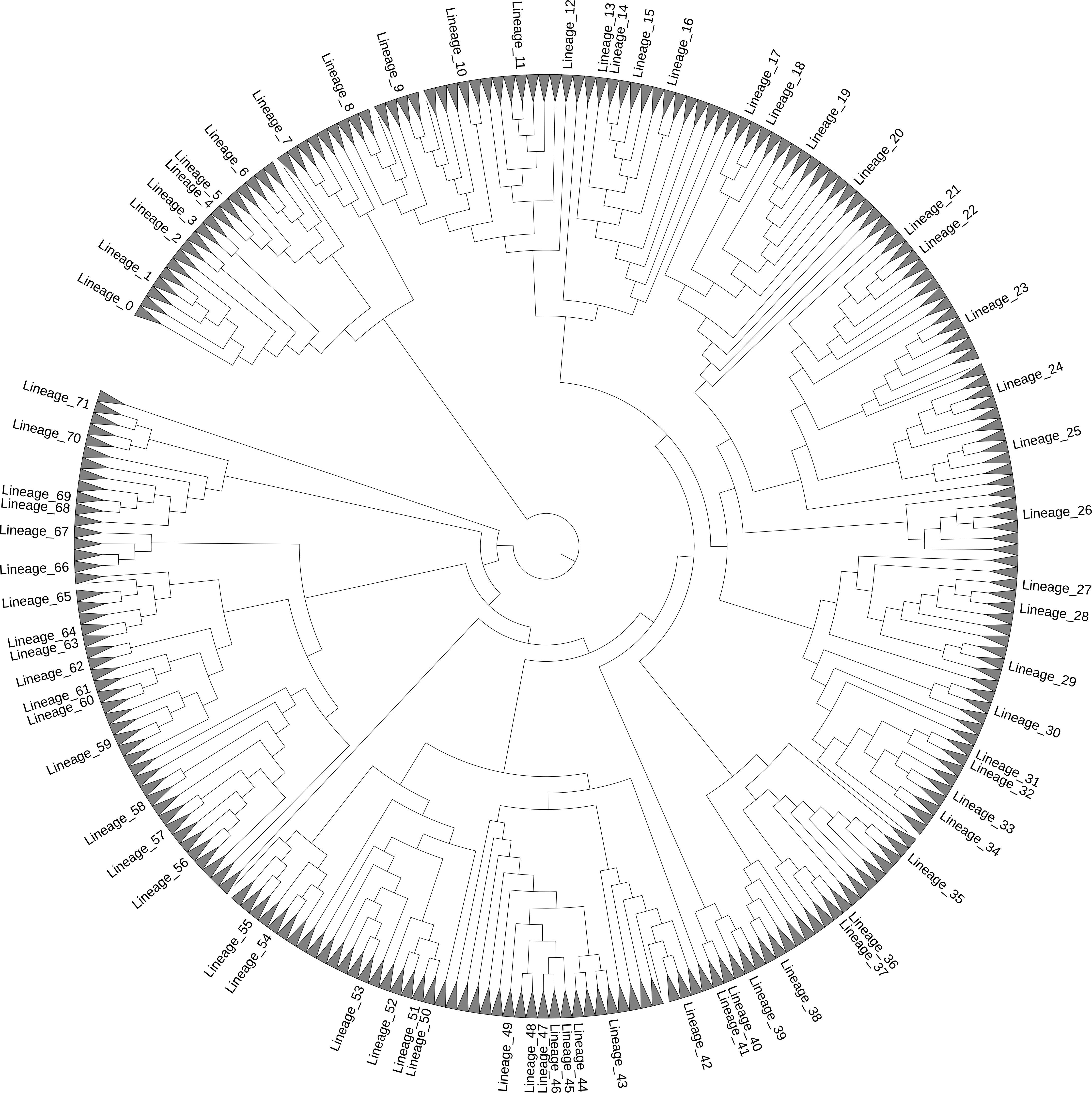

### Supplementary file 9

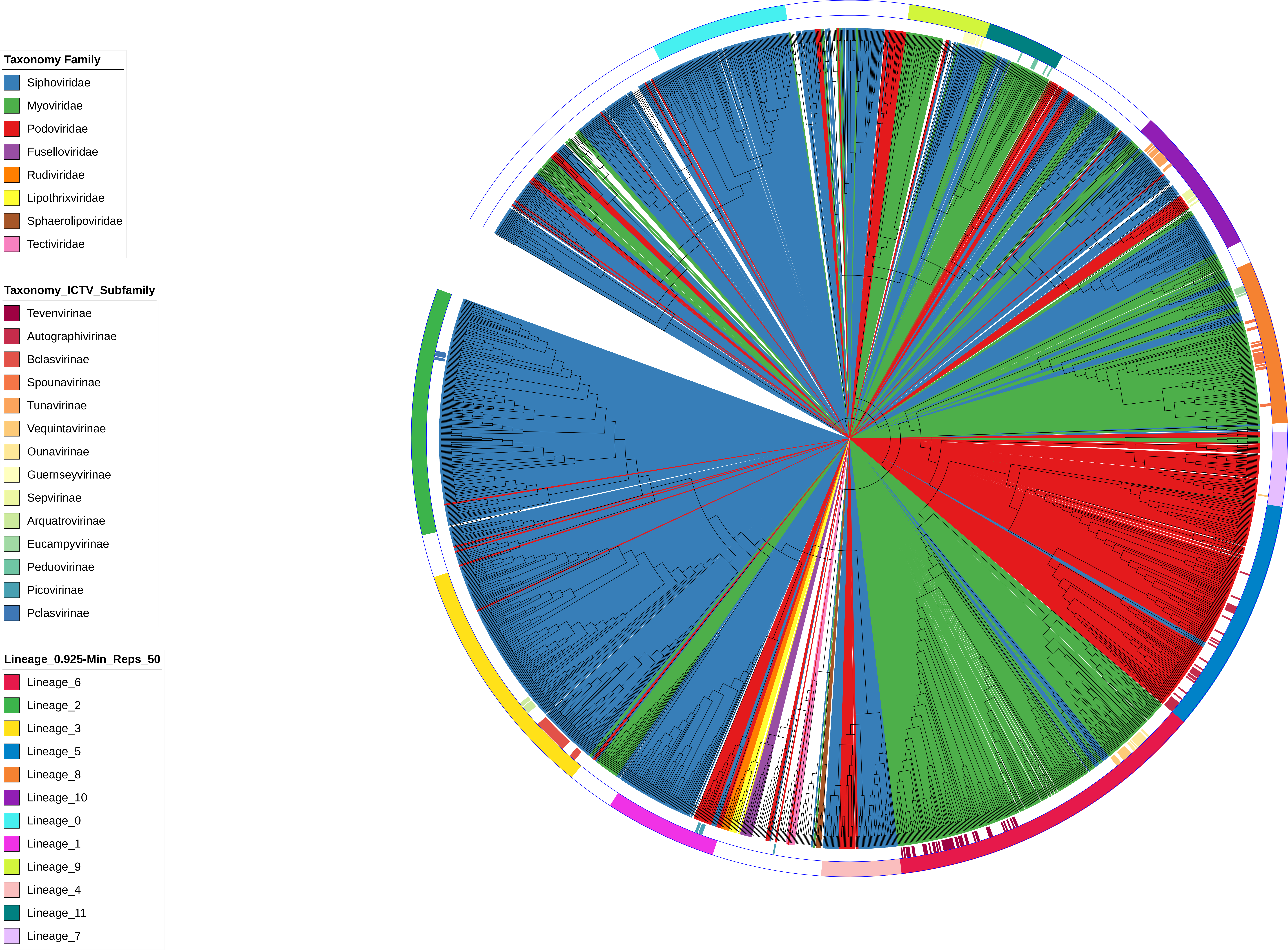

### Supplementary file 10

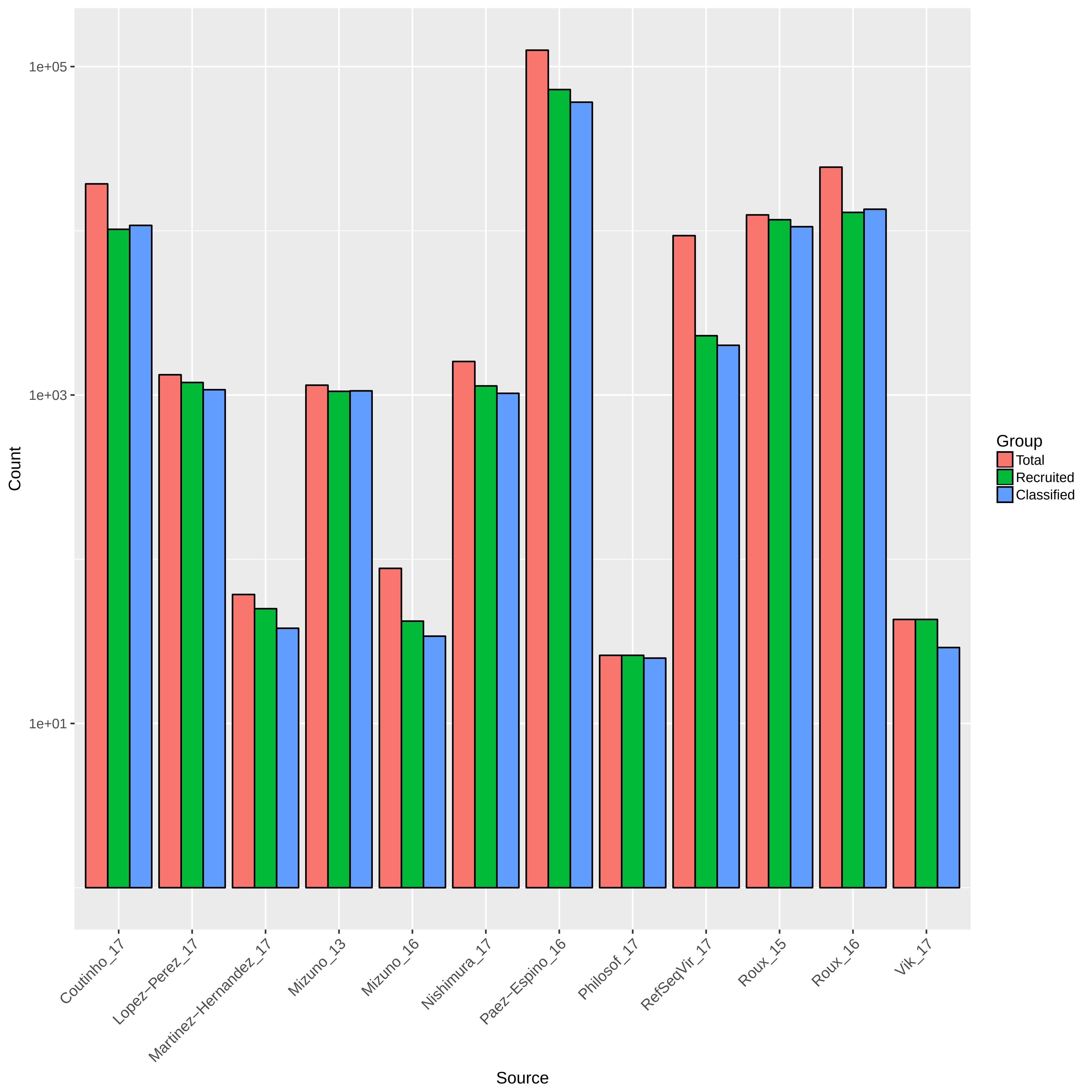

### Supplementary file 11

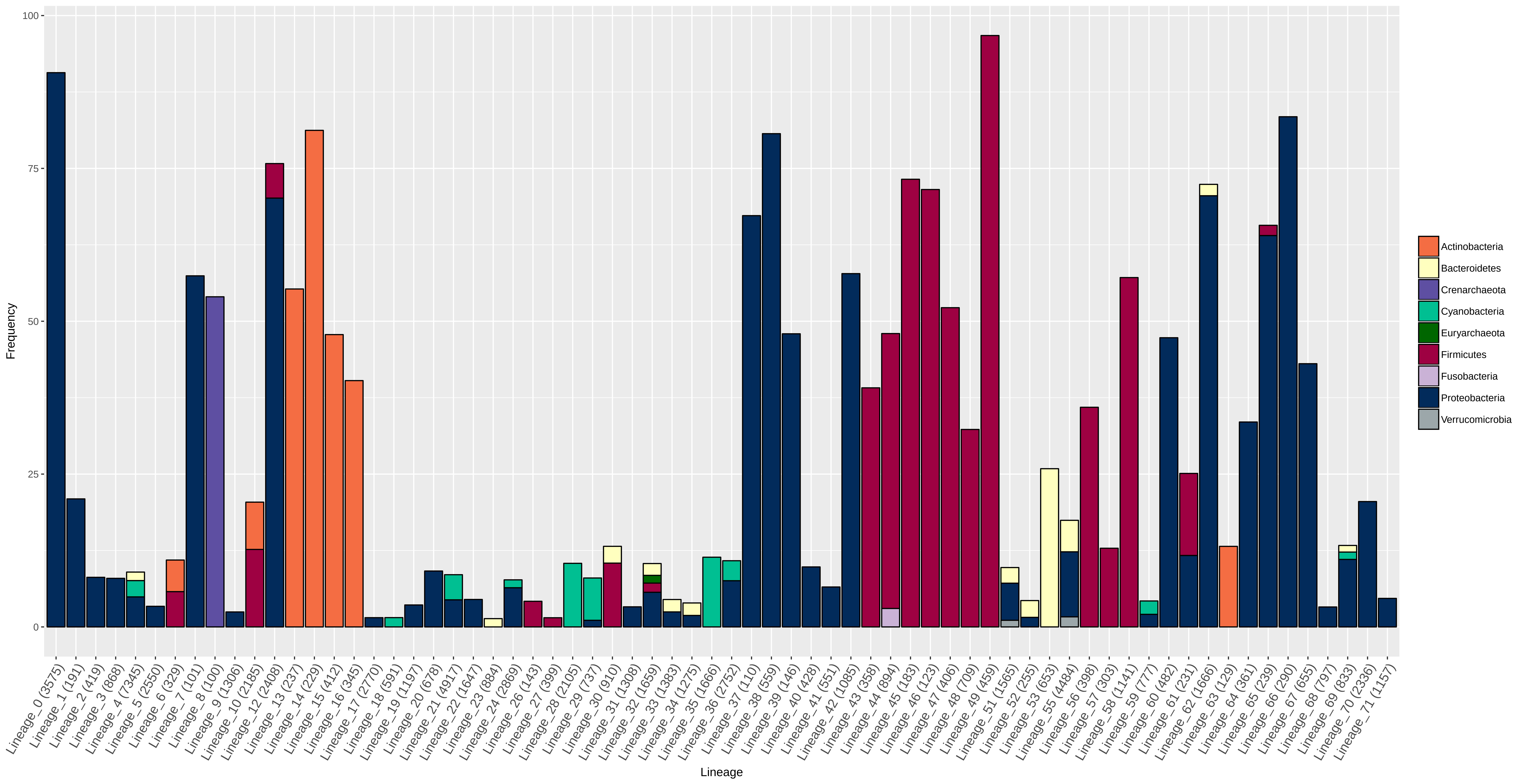

### Supplementary file 12

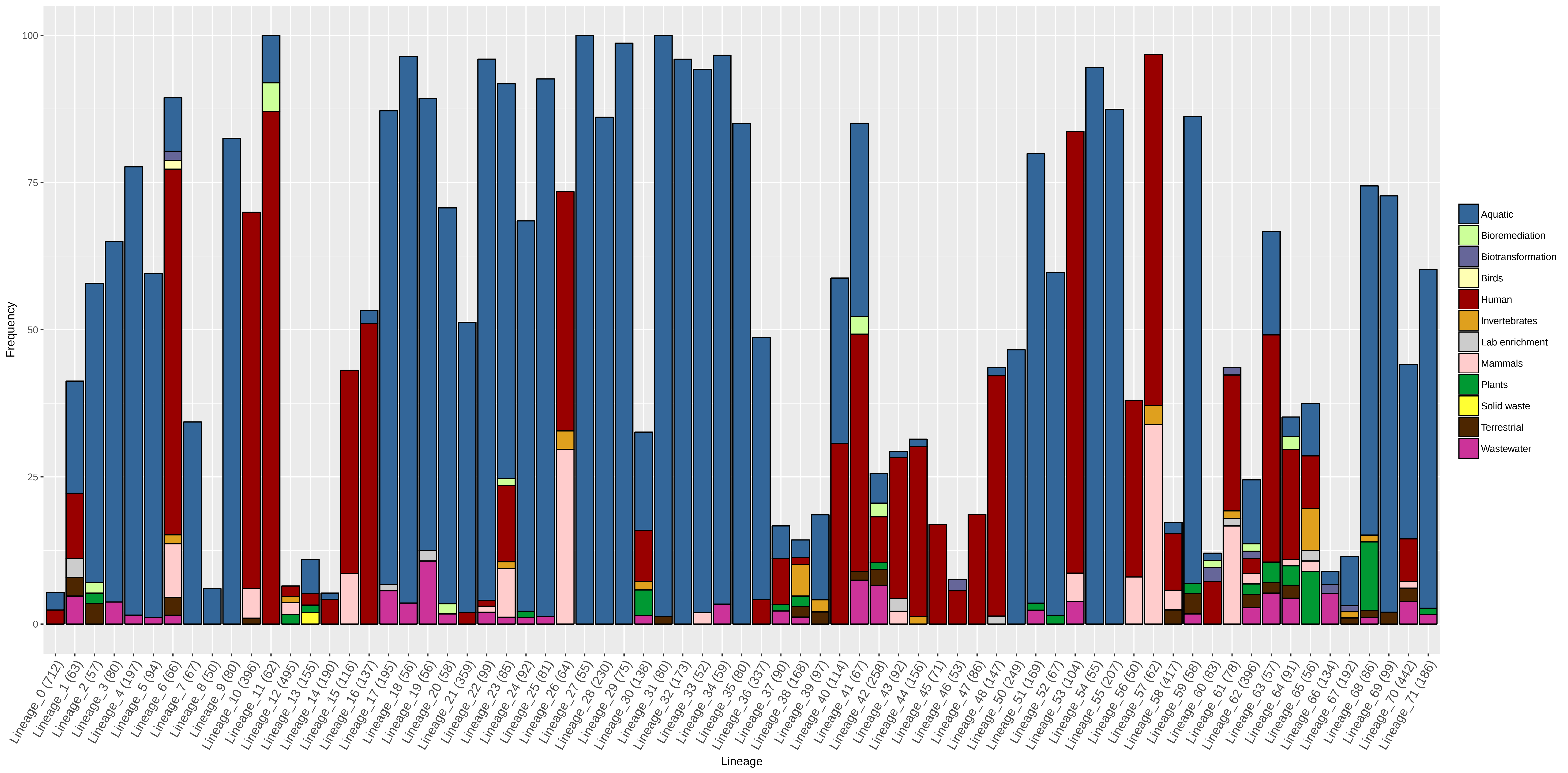
